# Node-degree aware edge sampling mitigates inflated classification performance in biomedical graph representation learning

**DOI:** 10.1101/2022.11.21.517376

**Authors:** Luca Cappelletti, Lauren Rekerle, Tommaso Fontana, Peter Hansen, Elena Casiraghi, Vida Ravanmehr, Christopher J Mungall, Jeremy Yang, Leonard Spranger, Guy Karlebach, J. Harry Caufield, Leigh Carmody, Ben Coleman, Tudor Oprea, Justin Reese, Giorgio Valentini, Peter N Robinson

## Abstract

Graph representation learning is a family of related approaches that learn low-dimensional vector representations of nodes and other graph elements called embeddings. Embeddings approximate characteristics of the graph and can be used for a variety of machine-learning tasks such as novel edge prediction. For many biomedical applications, partial knowledge exists about positive edges that represent relationships between pairs of entities, but little to no knowledge is available about negative edges that represent the explicit lack of a relationship between two nodes. For this reason, classification procedures are forced to assume that the vast majority of unlabeled edges are negative. Existing approaches to sampling negative edges for training and evaluating classifiers do so by uniformly sampling pairs of nodes. We show here that this sampling strategy typically leads to sets of positive and negative edges with imbalanced edge degree distributions. Using representative homogeneous and heterogeneous biomedical knowledge graphs, we show that this strategy artificially inflates measured classification performance. We present a degree-aware node sampling approach for sampling negative edge examples that mitigates this effect and is simple to implement.

## Introduction

Many problems in biology and medicine stand to benefit from machine learning (ML) approaches [1]. Biomedical data often are comprised of entities from multiple different classes that are interconnected by different types of relation. Therefore, biological data are often represented computationally as knowledge graphs (KG), semantic networks that encode entities as nodes and relations between entities as edges. Typical ML tasks that leverage KGs involve node (entity) classification and prediction of novel relations between entities (edge prediction) [2, 3]. Graph machine learning methods have been applied to numerous biomedical classification tasks including protein function prediction, protein–protein interaction prediction and in silico drug discovery [4]. Despite the great promise of ML in medicine, to date very few ML algorithms have contributed meaningfully to clinical care [5]. One reason for this might be that published models not infrequently display methodological flaws or underlying biases [6, 7, 8, 9]. It is therefore essential to understand and ideally to mitigate sources of bias and error in ML in order to develop robust and accurate algorithms.

Graph representation learning (GRL) is a form of graph machine learning that applies various strategies to convert nodes, edges, or graphs into low-dimensional vectors called “embeddings” that preserve graph structural information and properties [10]. Graph embeddings can be used to address downstream prediction tasks [11]. In this work, we focus on random-walk based GRL methods that optimize node embeddings such that nodes have similar embeddings if they tend to co-occur on short random walks over the graph [12, 3]. Numerous approaches have been developed to generate random walks that reflect different aspects of network structure, including DeepWalk, node2vec, and LINE [13, 14, 15] as well as approaches to assess co-occurrences of nodes in the random walks, such as Skip-Gram, Continuous Bag of Words, and GloVe [16, 17].

Recently, it was shown that topological imbalances in biomedical KGs can result in densely-connected entities (i.e., high-degree nodes) being highly ranked no matter the context, suggesting that embedding models may be more influenced by node degree rather than any biological information encoded within the relations [18]. This has been a well known property of any guilt by association analysis using biomedical knowledge for at least a decade [19]. We show that the method by which negative edges are sampled for ML can be responsible for this effect and present an approach to mitigating the effect by node-degree aware sampling. We demonstrate our approach using a homogeneous KG of protein-protein associations (PPAs) derived from the STRING database [20] and a heterogeneous graph in which synthetic lethal interactions [21, 22, 23] are integrated into the STRING PPA graph.

## Results

Here, we investigate the influence of negative-edge sampling on the measured performance of random-walk-based shallow graph representation learning (RW-GRL). In this framework, edge prediction involves the generation of low-dimensional vector representations (embeddings) of nodes based on the co-occurrence of pairs of nodes in random walks on the graph. Following this, edges are represented by transforming pairs of nodes by Hadamard transformation or related methods to obtain vectorial representations of edges [14]. Finally, classification methods such as random forest or neural networks are applied to the edge representations. For most biomedical networks, we have knowledge about a subset of the positive edges of interest (positive examples) but generally do not have information about explicitly excluded edges (negative examples). For this reason, most approaches to RW-GRL choose negative edges at random from edges that are not known to be positive under the assumption that the vast majority of unlabeled edges are negative.

### Sampling of negative examples in RW-GRL

There are three phases in RW-GRL that require sampling of negative examples: (i) During training of the embedding model, negative samples are chosen as the out-of-context nodes in models such as SkipGram or CBOW [16], or non-existent edges are sampled in models such as First-order LINE [15]. This first step is not required in some models such as GloVe [17] and HOPE [24]; (ii) During training of the classifier model for edge-prediction when the task is modeled as a binary prediction; and (iii) Finally, the evaluation step requires negative sampling to measure the generalization performance of the edge-prediction model. A common approach for obtaining negative examples samples the source and destination nodes from a uniform distribution. However, biomedical KGs are generally characterized by a non-uniform node-degree distribution [25], implying that edges sampled using the uniform sampling approach produce a graph with a degree distribution that can differ substantially from that of the existing edges.

Much previous work has been invested in understanding the effect of negative sampling in the first phase (training the embedding model). For instance, the node2vec algorithm scans over series of nodes encountered in random walks and attempts to predict nearby nodes (i.e., inside some context window) on the basis of a Skip-gram objective function. However, the per-node partition function is expensive to compute for large networks since it involves every node of the graph, and so node2vec approximates it using negative sampling, whereby *k* negative nodes are sampled for each positive node according to the unigram distribution U (w) raised to the 3/4rd power [14, 16]. A number of methods have been developed to improve the selection of negative examples for creating embeddings [26, 27] but there seems to be no single method that performs best for all datasets [28, 29]. In contrast, in the current work, we examine the effect of negative sampling methods on the evaluation phase.

### Classifiers can achieve an apparently good performance based solely on node-degree in KGs with unbalanced node distribution

To investigate this, we examined a homogeneous graph of protein-protein-associations (PPAs) derived from the STRING resource [20] and a heterogeneous graph consisting of the STRING-derived PPAs together with 2445 synthetic lethal interactions derived from the literature (SLDB/STRING; Methods). The classification task in the STRING graph was to predict novel PPAs, while the classification task in the heterogeneous SLDB/STRING graph was to predict novel synthetic lethal interactions.

Both graphs displayed a skewed node distribution. For instance, the mean degree of the top 20 most highly connected nodes in the STRING graph was 499.4, while mean degree in entire graph was only 30.5. 9875 of 252802 edges involved one of the top 20 nodes (3.9%). For the SLDB subgraph of the SLDB/STRING graph, the mean degree of the top 20 nodes was 105.9, compared to a mean degree in the entire graph of 2.8. 2081 of the 2445 edges (85.1%) in the largest component of the SLDB graph involved at least one of the top 20 nodes (Supplemental Figures S1-S4).

For each edge, we formed a two-dimension integer vector with the degree of each of the nodes that made up the edge. Using uniform node sampling, we then trained a perceptron to classify positive and negative edges. We present results of classification in terms of the Matthews correlation coefficient (MCC), which ranges from −1 for perfect misclassification to +1for perfect classification, while MCC=0 is the expected value for a random classifier [30].

We performed ten-fold cross validation with training size of 0.75. An equal number of positive and negative examples was chosen. For the STRING graph, we obtained an MCC of 0.91 ± 0.01 (train) and 0.87 ± 0.01(test). For the SLDB/STRING graph, the results were 0.905 ± 0.007 (train) 0.879 ± 0.010 (test). These results show that a model predicting the existence of an edge solely on the basis of the node degree of the source and destination nodes may obtain an apparently good performance, whether or not node degree reflects the biological functions being investigated, suggesting the possibility that bias in the construction or evaluation of the model could be artificially inflating the measured performance of the classifier.

### Two methods for sampling negative examples

A common approach for obtaining negative examples samples the source and destination nodes from a uniform distribution that randomly chooses an integer between 1 and |*V* | corresponding to the nodes of the graph (Figure 2A). We reasoned that this sampling strategy, which we term *edge sampling by uniform node sampling* (UNS) will produce negative examples whose node degree approximates the degree distribution of the entire KG but may differ from the node degree distribution of the positive examples in many relevant biomedical KGs.

**Figure 1:**
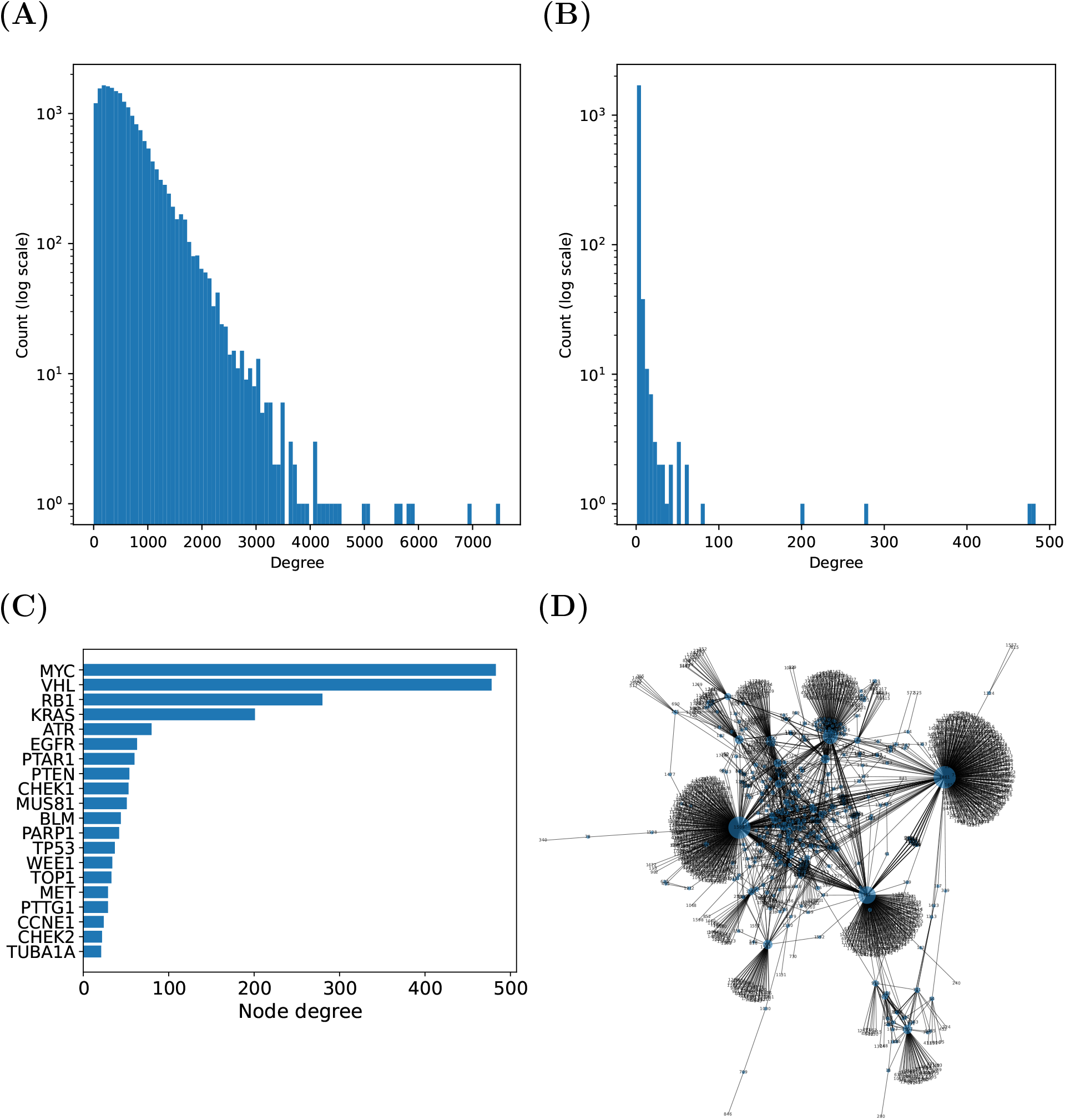
**STRING graph** Topology of the STRING and SLDB/STRING graphs. **A)** Degree distribution of the STRING protein-protein association graph. **B)** Degree distribution of the SLDB (synthetic lethality interaction database) component of the SLDB/STRING graph. **C)** The 20 highest-degree nodes of the SLDB network (which is a component of the SLDB/STRING graph). The distribution of node degrees among the 20 most densely connected nodes of the SLI network. **D)** Visualization of the SLDB network. Only the largest connected component is shown. Nodes with degree of 2 or more are shown in blue. Inspection of the SLDB graph showed it to have a highly skewed degree distribution with a few hubs, i.e., nodes that are highly connected to other nodes in the network, and many nodes with few connections. See Supplemental Figure 1 for an analysis of the degree distribution of this graph.

**Figure 2:**
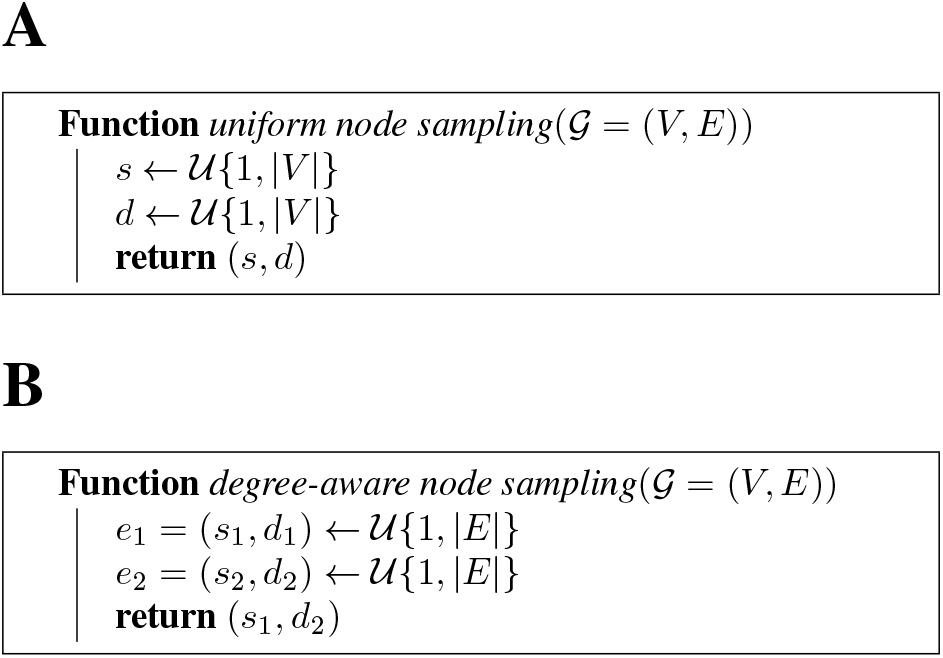
Pseudocode for edge sampling strategies. Both strategies sample two nodes and return the edge that connects the two nodes, but the procedure used for sampling the nodes differs. **A**. **uniform node sampling (UNS)**. The standard method for sampling samples two nodes uniformly to create a “random” edge for negative examples. **B**. **degree-aware node sampling (DANS)**. The method presented here instead samples two edges uniformly to create a random edge from the source node of the first edge and the destination node of the second edge.

We therefore developed a different sampling mechanism that assigns a number of negative edges to each node proportional to its node degree. We term this method *edge sampling by degree-aware node sampling* (DANS). In this approach, we randomly sample two edges *e*_1_ = (*s*_1_, *d*_2_), *e*_2_ = (*s*_2_, *d*_2_) from 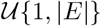, and build a new negative edge by connecting the source node of *e*_1_ and the destination node of *e*_2_ (Figure 2B).

In real-world graphs, there is a minimal likelihood of collisions between existent and non-existent edges using either the UNS or the DANS sampling strategy that can be trivially addressed by repeating the sampling.

### UNS- and DANS-based node-degree-based classification

We then repeated the above experiment in which we trained a perceptron to classify edges based on a two-dimensional vector with the degrees of the nodes making up each edge. We compared the measured performance of this classificer using the UNS-based and DANS-based negative edge sampling procedures in the STRING and SLDB/STRING graphs. For the STRING graph, while standard UNS-based sampling achieved a MCC of over 0.5, DANS-based sampling achieved a MCC near zero, which corresponds to random guessing. The MCC for the SLDB/STRING graph was over 0.85 for UNS sampling but less than 0.4 for DANS-based sampling in the test phase (Figure 3; see also Supplemental Tables S2 and S3 for corresponding evaluations using the area under the receiver operating characteristics curve (AUROC), area under the precision recall curve (AUPRC), and the F1 score).

**Figure 3:**
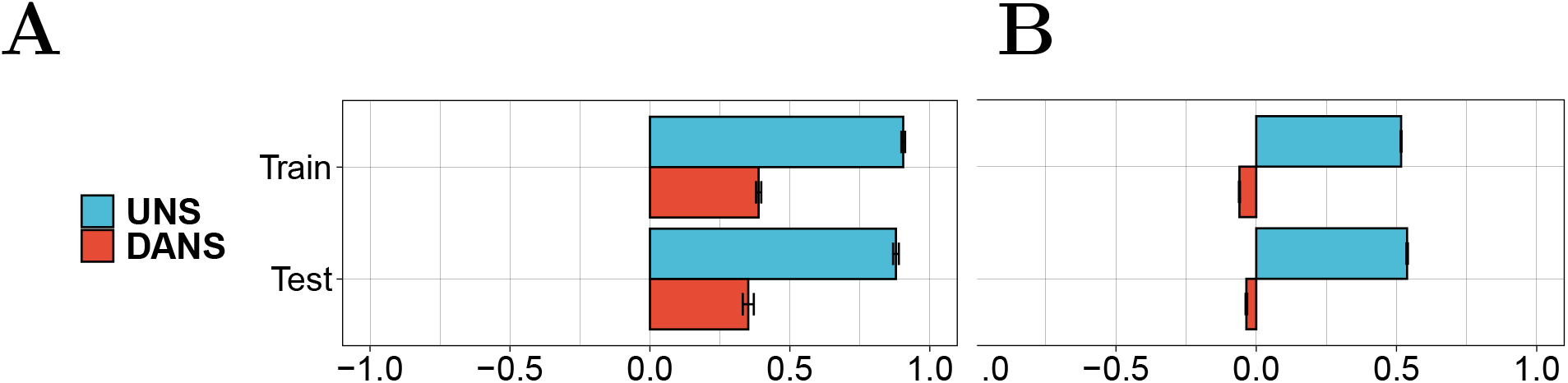
Matthews correlation coefficient (MCC) analysis for classification of (A) SLDB/STRING edges and (B) STRING edges. In both edge prediction problems, the measured performance with uniform sampling was substantially higher than with node-based sampling.

Both classification models were created identically with the sole exception of the method for choosing negative examples. This suggests that classification performance measured for the UNS sampling approach may be inflated. In the STRING graph, the performance of the classification using DANS sampling was close to that of random guessing, which is what we would expect if the biological signal is not related to node degree. For the SLDB/STRING graph, the measured performance of classification using DANS sampling was substantially less than with UNS sampling. The difference likely reflects the fact that the degree distribution of the SLDB graph is more highly skewed than that of the STRING graph (Figure 1). Both experiments suggest that at least part of the signal obtained by the classifier using UNS sampling is spurious.

### The influence of negative sampling strategies on RW-GRL

We then asked if a similar effect pertains to RW-GRL. We applied nine different embedding approaches followed by perceptron-based classification of edges in both the STRING and the SLDB/STRING graphs (Methods).

For SLDB/STRING, we tested the classification performance for the prediction of synthetic lethal interaction edges, one of the two types of edges in the graph in addition to PPA edges. For instance, Walklets SkipGram displayed an MCC of 0.74 for UNS sampling and 0.32 for DANS sampling (test). With the exception of one model (DeepWalk GloVe), the measured results of models built with UNS sampling were higher than those obtained with the same models using DANS sampling, with the difference in MCC ranging from 0.007 for Walklets GloVe to 0.57 for Second-order LINE (Figure 4; see also Supplemental Table S4 for AUROC, AUPRC, and F1 score analysis). For each comparison the only difference was in the way the negative examples were selected, suggesting again classification performance measured for the UNS approach may be inflated for RW-GRL classifiers.

**Figure 4:**
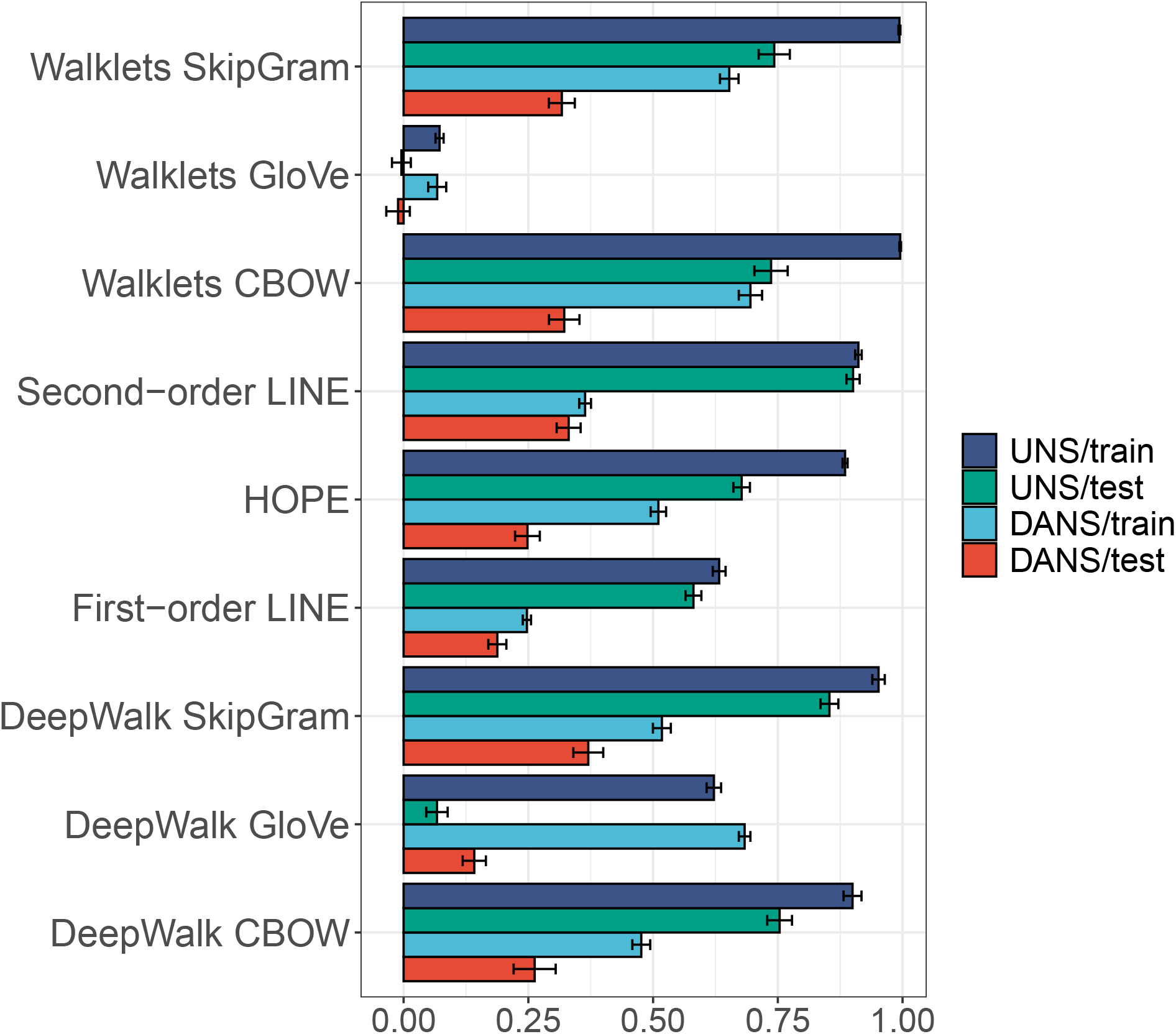
Matthews correlation coefficient for the nine RW-GRL methods applied to edge prediction in the heterogeneous SLDB/STRING graph.

One model, DeepWalk GloVe, showed marginally better performance with DANS sampling than with UNS sampling. This model was the eighth worst performing model as assessed by the train and test results (Supplemental Tables S4). We do not have an explanation for this observation, but note that it underlines the importance of performing in-depth diagnostics of RW-GRL models.

We then repeated the experiments for the homogeneous STRING network, for which we tested the classification performance for the prediction of PPA edges. The results again showed that classification performed with uniform sampling of negative edges showed higher apparent classification performance than the same experiments using node-base negative edge sampling (Figure 5 and Supplemental Table S5).

**Figure 5:**
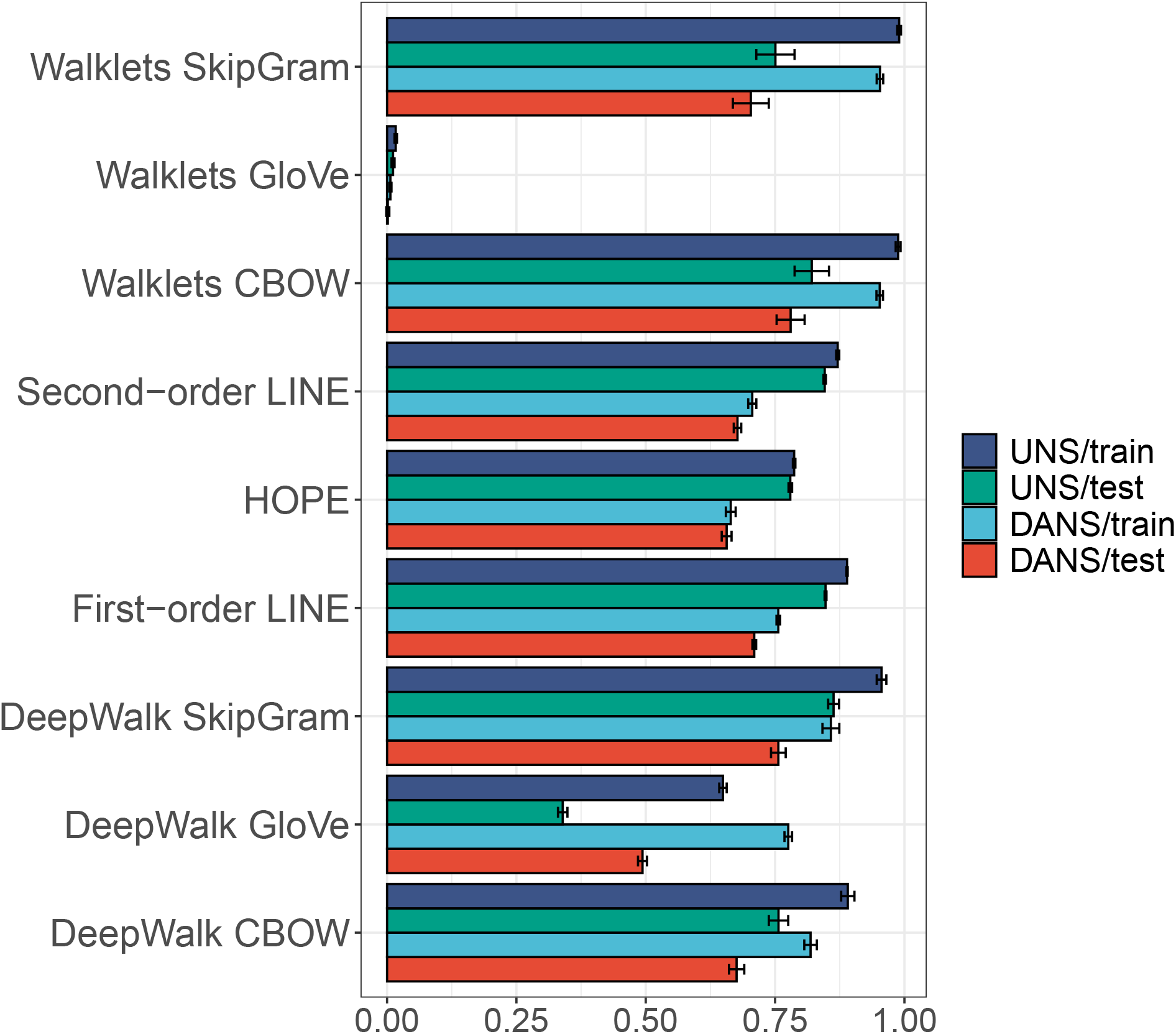
Matthews correlation coefficient for the nine RW-GRL methods applied to edge prediction in the homogeneous STRING graph.

The difference in the influence of negative sampling strategies on the SLDB/STRING and STRING graphs is related to the different node distributions or other differences in graph structure. Indeed, the SLDB/STRING graph shows a degree distribution that is approximately scale-free, with few nodes of very high degree and many low-degree nodes (Supplemental Figure S1). The STRING graph shows a skewed but more even degree distribution (Figure 1 and Supplemental Table S1).

## Discussion

Constructing generalizable graph models of biomedical domains is challenging because most systems of scientific interest have multiple different classes of nodes and relations, and in many cases, our knowledge is so incomplete that comprehensive gold-standard data sets for training ML classifiers do not exist. An ML algorithm is said to be biased if its results are systematically wrong due to incorrect assumptions of the ML process. Biases can inflate the measured prediction performance of algorithms [31]. In this work we have explored the relationship between sampling of negative examples and measured performance of shallow graph representation learners. Our results demonstrate that if the positive and negative samples have a different node degree distribution, then sampling strategies that do not take node degree into account can inflate the measured performance of classification algorithms because the choice of negative samples is biased to true negatives, hence causing a positive bias. Skewed node degree distribution (a specific kind of topological imbalance) has been increasingly recognized as a source of bias for bioinformatic algorithms [18]. Our results demonstrate a factor that contributes to this bias and provide a simple strategy for mitigation.

The standard method of uniform sampling randomly chooses pairs of nodes without taking node degree into account. This approach is biased in cross-validation settings, because the positive training examples will have higher node degrees that the negative examples. Therefore, if we sample edges by randomly sampling pairs of nodes, we will be more likely to choose negative edges and will thus give the classifier an advantage. If on the other hand, we choose negative edges by randomly sampling edges, there is no bias according to node degree. To our knowledge, there is no existing software to systematically investigate biases of this type in RW-GRL experiments. Our review of other software packages for RW-GRL revealed that the uniform node sampling procedure is used during training by the gensim package [32] that is widely used as to develop algorithms to perform embedding such as node2vec [14]. The methods used for sampling negative examples are rarely described in detail in the methods of many publications on the topic.

In RW-GRL, there are three phases in which negative examples are sampling. The bias that we describe here is primarily related to sampling in the third phase, which evaluates the predictions of novel links. In preliminary experiments, we did not find a signal across all nine tested models when we limited DANS sampling to just the first stage (construction of the embeddings) or the second stage (data not shown). However, for specific models and specific learning tasks, it may be useful to test the effects of the two sampling procedures on each of the three stages separately.

In general, negative samples should be meaningful for the classification task at hand, and inappropriate sampling mechanisms may lead to biased evaluation. Negative samples that differ substantially from the positive samples in one or more characteristics can lead to over-optimistic evaluations. Conversely, negative samples that are too similar to (or even collide with) the positive samples may lead to overly poor evaluations. We hypothesized that a method for sampling negative edges that samples according to node degree would generate negative examples whose degree distribution would more closely resemble that of the positive examples. To do so, we uniformly sample two edges and choose the source node of the first edge and the destination node of the second edge to choose edges for the negative set (Figure 2). For both approaches, if an edge is chosen that is already in the positive or negative sets, we simply repeat the procedure.

We recommend that practitioners of RW-GRL carefully evaluate the node degree distribution of samples chosen for the negative and positive examples. Results of training with the standard uniform node selection schema and the node-degree selection approach presented here should be compared.

We note that many biological networks do show a highly skewed (if not necessarily scale-free) node distribution. For instance, the most densely connected genes in the SLDB graph represent genes with central roles in oncogenesis, such as *MYC, VHL*, and *RB1* (Supplemental Table S2). This may represent biological reality or may reflect an experimental ascertainment bias that is not mitigated by the sampling scheme we present here. Detailed case-by-case assessment of the data will be needed to understand how to apply graph ML and interpret the results in light of a potential node degree bias.

## Methods

### Protein-Protein Associations

The STRING database is a comprehensive relational database of protein-protein associations [20]. STRING (version 11.0) data for *H. sapiens* was used corresponding to `9606.protein.links.v11.5.txt.gz`. Associations were filtered to retain only those with a score of at least 700, and duplicate edges between node pairs were removed.

### Synthetic lethal interaction data

We first analyzed data available in the supplemental data provided with ISLE [33] and files available from SynLethDB resource [34] by comparing the curation with the original publications. To create the synthetic lethal interaction database (SLDB) resource, We manually reviewed publications cited in these resources and additional publications. The curated SLIs are available at https://github.com/monarch-initiative/syntheticLethalityNetwork. The tab-separated file (TSV) includes information about each pair of genes, the perturbations used for each gene, the assays used to measure SL, a Cellosaurus id [35] (if applicable), and the PubMed identifier. To add additional information to SLDB, we integrated it with the STRING protein-protein interaction network (SLDB/STRING) by using the the nodes (genes) in SLDB to also represent the proteins in STRING that the genes themselves encode.

For the experiments described in this work, we imported the SLDB resource directly using a utility function of GRAPE [36].

### Shallow Graph Representation Learning

The shallow graph representation learning experiments described here were performed using the GRAPE library for fast and scalable Graph Processing and Embedding, version 0.1.28. GRAPE provides a comprehensive library of Graph Representation Learning and inference models implemented in Rust with a Python interface [36].

Nine node embedding methods were used including neural network-based methods and matrix factorization methods. The former include methods based on edge sampling such as first and second-order LINE [15], which trains a neural network with either one layer (first-order) or two layers (second-order) to predict whether a given tuple of nodes defines an existing edge. Additionally, neural network-based methods included methods based on random walk sampling: we have considered two random walk sampling mechanisms, namely DeepWalk [13], which samples first-order random walks, and Walklets [37], which samples first-order random walks and, for a given central node *v*, at each *i*-th sampling iteration, skips *i* nodes around the central node *v*. We employed the training samples obtained with both DeepWalk and Walklets to train three different embedding models. First, the GloVe [17] model, which trains a second-order siamese neural network to predict the log-normalized co-occurrence of two nodes within a sliding window over the random walk samples. Second, the CBOW [38] model, trains a second-order siamese neural network to predict the central node of a random walk window given the remainder contextual nodes. The third and final considered model is SkipGram [38], which analogously to CBOW trains a second-order siamese neural network to predict the contextual nodes given the central node. In all of the considered neural network models, the resulting node embedding matrix coincides with the (trained) weight matrix of the first hidden layer. Finally, we have employed two matrix factorization methods, namely HOPE [24], which starts by computing a node-proximity matrix, where the proximity between two nodes may be defined in different ways, in our case by using the number of common neighbors. Then HOPE computes the singular vectors corresponding to the *k* most significant singular values of the proximity matrix and uses the left and right product of the singular values with the singular vectors as the embeddings of the source and destination nodes. The second matrix factorization method we considered is NetMF [39], which given a window-size, first computes a sparse log co-occurrence matrix by using first-order random walks and then proceeds to compute the singular vectors corresponding to the *k* largest singular values.

Each of the nine algorithms was used with GRAPE default parameters. Details are provided in Table 1.

**Table 1:** Parameters for the learning models used in this project. Each of the models shown here was run using uniform and node-based sampling of negative examples. In addition to the parameters shown in the table, some parameters are only relevant to a subset of models: `learning rate decay` was set to 0.9 (for all models except HOPE and NetMF). `avoid false negatives` was set to false for First-order LINE and Second-order LINE. `alpha` was set to 0.75 for DeepWalk GloVe and Walklets GloVe. The metric used for HOPE was `neighbors intersection size`. The `number of negative samples` was set to 10 for DeepWalk CBOW, DeepWalk SkipGram, Walklets CBOW, and Walklets SkipGram. `Iterations` was set to 100 for all models except First-order LINE, Second-order LINE, and HOPE.

| Model | Epochs | Learning rate | Walk length | Window size | Max neighbors |
| --- | --- | --- | --- | --- | --- |
| DeepWalk CBOW | 30 | 0.010 | 128 | 5 | 100 |
| DeepWalk GloVe | 30 | 0.001 | 128 | 5 | 100 |
| DeepWalk SkipGram | 30 | 0.010 | 128 | 5 | 100 |
| First-order LINE | 100 | 0.050 | n/a | n/a | n/a |
| HOPE | n/a | n/a | n/a | n/a | n/a |
| Second-order LINE | 100 | 0.050 | n/a | n/a | n/a |
| Walklets CBOW | 30 | 0.010 | 128 | 4 | 100 |
| Walklets GloVe | 30 | 0.001 | 128 | 4 | 100 |
| Walklets SkipGram | 30 | 0.010 | 128 | 4 | 100 |

Edge embeddings were formed from the embeddings of the corresponding pair of nodes (*u,v*) by the binary Hadamard (⊡) operator, defined as the elementwise product of both vectors, i.e.,

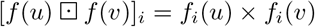

Edge prediction was performed by a Perceptron model.

## Code availability

The scripts used to generate the results described in this manuscript are availabile together with several illustrative Jupyter notebooks at https://github.com/monarch-initiative/negativeExampleSelection under the GNU General Public License version 3.

## Funding

This work was funded by NIH [U01-CA239108-02]; Additional support was received from NCI grant U24-CA224067.

## Notes

### Competing Interest Statement

The authors have declared no competing interest.

