## Supplement for "Node-degree aware edge sampling mitigates inflated classification performance in biomedical graph representation learning"

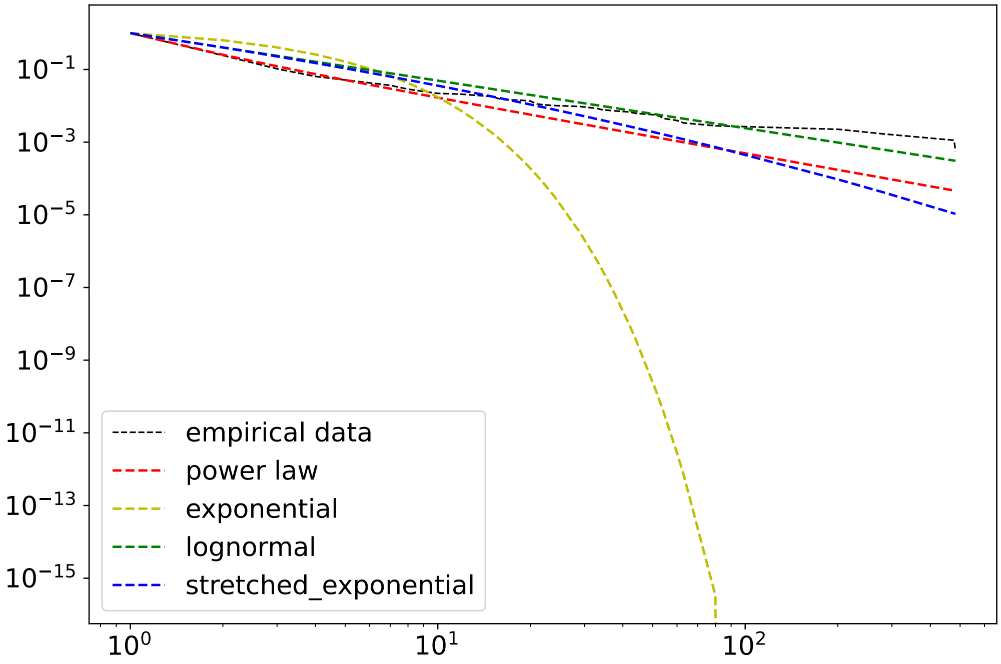


**Figure S1: Analysis of the degree distribution of the SLDB graph.** The observed distribution was consistent with a power-law distribution with $\alpha = 2.5$. No other tested distribution provided a significantly better fit. The largest connected component of the graph was analyzed with the the powerlaw Python package [1].

| Characteristic | SLDB | STRING | SLDB/STRING |
|---|---|---|---|
| connected components | 6 | 98 | 1 |
| size of smallest component | 2 | 2 | 16715 |
| size of largest component | 1765 | 16582 | 16715 |
| nodes | 1775 | 16812 | 16715 |
| edges | 2445 | 252953 | 255253 |
| diameter | 9.0 | 14.0 | 14.0 |
| clustering coefficient | 58.4 | 5617.6 | 5588.4 |
| density | 0.0016 | 0.0018 | 0.0017 |
| mean node degree | 2.8 | 30.5 | 30.5 |
| median node degree | 1 | 13 | 13 |
| transitivity | 0.0034 | 0.31 | 0.30 |

**Table S1: Network characteristics of the SLDB, STRING, and SLDB/STRING (composite) graphs**. Network characteristics were calculated with the GRAPE library [2].

| Model | Evaluation | neg. select | F1 | AUROC | AUPRC | MCC |
|---|---|---|---|---|---|---|
| Degree | train | Node-degree | $0.563 \pm 0.002$ | $0.491 \pm 0.001$ | $0.517 \pm 0.001$ | $-0.060 \pm 0.002$ |
| Degree | test | Node-degree | $0.576 \pm 0.002$ | $0.508 \pm 0.002$ | $0.528 \pm 0.002$ | $-0.035 \pm 0.003$ |
| Degree | train | Uniform | $0.735 \pm 0.003$ | $0.841 \pm 0.000$ | $0.844 \pm 0.001$ | $0.516 \pm 0.002$ |
| Degree | test | Uniform | $0.750 \pm 0.003$ | $0.856 \pm 0.001$ | $0.855 \pm 0.001$ | $0.538 \pm 0.003$ |

**Table S2:** STRING: Node degree bias analysis. For each edge, we formed a two-dimension integer vector with the degree of each of the nodes that made up the edge. Using uniform node sampling, we then trained a perceptron to classify positive and negative edges. F1: F1 score (harmonic mean of the precision and recall); AUROC: area under the receiver operating characteristics; AUPRC: area under the precision recall curve. Results are shown for train and test phases using the two different negative example selection strategies investigated in this work: uniform node sampling, and node-degree sampling.

| Model | Evaluation | neg. select | F1 | AUROC | AUPRC | MCC |
|---|---|---|---|---|---|---|
| Degree | test | Node-degree | $0.719 \pm 0.007$ | $0.644 \pm 0.011$ | $0.548 \pm 0.008$ | $0.351 \pm 0.020$ |
| Degree | train | Node-degree | $0.734 \pm 0.003$ | $0.665 \pm 0.004$ | $0.562 \pm 0.003$ | $0.388 \pm 0.009$ |
| Degree | test | Uniform | $0.932 \pm 0.007$ | $0.995 \pm 0.002$ | $0.994 \pm 0.003$ | $0.879 \pm 0.010$ |
| Degree | train | Uniform | $0.948 \pm 0.004$ | $0.999 \pm 0.001$ | $0.996 \pm 0.003$ | $0.905 \pm 0.007$ |

**Table S3:** SLDB/STRING: Node degree bias analysis. Explanations and abbreviations as in Table **S2**.

| Model | Evaluation | neg. select | F1 | AUROC | AUPRC | MCC |
|---|---|---|---|---|---|---|
| DeepWalk CBOW | test | Node-degree | $0.676 \pm 0.017$ | $0.635 \pm 0.024$ | $0.572 \pm 0.021$ | $0.263 \pm 0.042$ |
| DeepWalk CBOW | train | Node-degree | $0.765 \pm 0.006$ | $0.809 \pm 0.034$ | $0.744 \pm 0.042$ | $0.476 \pm 0.018$ |
| DeepWalk CBOW | test | Uniform | $0.861 \pm 0.018$ | $0.951 \pm 0.010$ | $0.939 \pm 0.015$ | $0.754 \pm 0.025$ |
| DeepWalk CBOW | train | Uniform | $0.950 \pm 0.009$ | $0.983 \pm 0.004$ | $0.960 \pm 0.009$ | $0.900 \pm 0.018$ |
| DeepWalk GloVe | test | Node-degree | $0.321 \pm 0.022$ | $0.584 \pm 0.012$ | $0.592 \pm 0.014$ | $0.142 \pm 0.023$ |
| DeepWalk GloVe | train | Node-degree | $0.830 \pm 0.006$ | $0.928 \pm 0.004$ | $0.923 \pm 0.006$ | $0.683 \pm 0.012$ |
| DeepWalk GloVe | test | Uniform | $0.308 \pm 0.021$ | $0.597 \pm 0.011$ | $0.561 \pm 0.015$ | $0.067 \pm 0.021$ |
| DeepWalk GloVe | train | Uniform | $0.805 \pm 0.008$ | $0.902 \pm 0.004$ | $0.887 \pm 0.005$ | $0.622 \pm 0.014$ |
| DeepWalk SkipGram | test | Node-degree | $0.725 \pm 0.011$ | $0.725 \pm 0.019$ | $0.657 \pm 0.019$ | $0.370 \pm 0.030$ |
| DeepWalk SkipGram | train | Node-degree | $0.779 \pm 0.007$ | $0.819 \pm 0.014$ | $0.740 \pm 0.018$ | $0.518 \pm 0.018$ |
| DeepWalk SkipGram | test | Uniform | $0.921 \pm 0.010$ | $0.968 \pm 0.007$ | $0.966 \pm 0.012$ | $0.853 \pm 0.018$ |
| DeepWalk SkipGram | train | Uniform | $0.976 \pm 0.006$ | $0.994 \pm 0.003$ | $0.982 \pm 0.009$ | $0.952 \pm 0.013$ |
| First-order LINE | test | Node-degree | $0.641 \pm 0.007$ | $0.631 \pm 0.013$ | $0.622 \pm 0.018$ | $0.188 \pm 0.018$ |
| First-order LINE | train | Node-degree | $0.671 \pm 0.003$ | $0.667 \pm 0.004$ | $0.629 \pm 0.006$ | $0.247 \pm 0.008$ |
| First-order LINE | test | Uniform | $0.775 \pm 0.009$ | $0.831 \pm 0.009$ | $0.873 \pm 0.008$ | $0.581 \pm 0.016$ |
| First-order LINE | train | Uniform | $0.810 \pm 0.006$ | $0.888 \pm 0.005$ | $0.904 \pm 0.004$ | $0.632 \pm 0.013$ |
| HOPE | test | Node-degree | $0.639 \pm 0.013$ | $0.659 \pm 0.019$ | $0.633 \pm 0.028$ | $0.248 \pm 0.025$ |
| HOPE | train | Node-degree | $0.778 \pm 0.006$ | $0.771 \pm 0.010$ | $0.690 \pm 0.032$ | $0.511 \pm 0.015$ |
| HOPE | test | Uniform | $0.790 \pm 0.014$ | $0.897 \pm 0.017$ | $0.926 \pm 0.010$ | $0.678 \pm 0.017$ |
| HOPE | train | Uniform | $0.939 \pm 0.003$ | $0.986 \pm 0.001$ | $0.981 \pm 0.005$ | $0.885 \pm 0.005$ |
| Second-order LINE | test | Node-degree | $0.718 \pm 0.007$ | $0.730 \pm 0.014$ | $0.691 \pm 0.018$ | $0.331 \pm 0.024$ |
| Second-order LINE | train | Node-degree | $0.727 \pm 0.004$ | $0.750 \pm 0.006$ | $0.709 \pm 0.010$ | $0.364 \pm 0.012$ |
| Second-order LINE | test | Uniform | $0.951 \pm 0.007$ | $0.986 \pm 0.003$ | $0.985 \pm 0.004$ | $0.901 \pm 0.013$ |
| Second-order LINE | train | Uniform | $0.956 \pm 0.003$ | $0.988 \pm 0.001$ | $0.987 \pm 0.002$ | $0.911 \pm 0.007$ |
| Walklets CBOW | test | Node-degree | $0.660 \pm 0.025$ | $0.671 \pm 0.026$ | $0.610 \pm 0.026$ | $0.322 \pm 0.031$ |
| Walklets CBOW | train | Node-degree | $0.852 \pm 0.011$ | $0.997 \pm 0.002$ | $0.993 \pm 0.004$ | $0.695 \pm 0.023$ |
| Walklets CBOW | test | Uniform | $0.827 \pm 0.028$ | $0.982 \pm 0.006$ | $0.981 \pm 0.004$ | $0.736 \pm 0.033$ |
| Walklets CBOW | train | Uniform | $0.998 \pm 0.001$ | $0.999 \pm 0.001$ | $0.999 \pm 0.002$ | $0.995 \pm 0.002$ |
| Walklets GloVe | test | Node-degree | $0.487 \pm 0.014$ | $0.501 \pm 0.010$ | $0.490 \pm 0.009$ | $-0.011 \pm 0.024$ |
| Walklets GloVe | train | Node-degree | $0.543 \pm 0.011$ | $0.546 \pm 0.008$ | $0.525 \pm 0.007$ | $0.067 \pm 0.018$ |
| Walklets GloVe | test | Uniform | $0.489 \pm 0.013$ | $0.505 \pm 0.008$ | $0.476 \pm 0.006$ | $-0.004 \pm 0.019$ |
| Walklets GloVe | train | Uniform | $0.545 \pm 0.008$ | $0.549 \pm 0.007$ | $0.507 \pm 0.006$ | $0.072 \pm 0.008$ |
| Walklets SkipGram | test | Node-degree | $0.672 \pm 0.017$ | $0.681 \pm 0.026$ | $0.632 \pm 0.028$ | $0.317 \pm 0.026$ |
| Walklets SkipGram | train | Node-degree | $0.833 \pm 0.008$ | $0.993 \pm 0.005$ | $0.987 \pm 0.009$ | $0.653 \pm 0.019$ |
| Walklets SkipGram | test | Uniform | $0.833 \pm 0.026$ | $0.958 \pm 0.010$ | $0.965 \pm 0.009$ | $0.743 \pm 0.031$ |
| Walklets SkipGram | train | Uniform | $0.997 \pm 0.001$ | $0.999 \pm 0.001$ | $0.998 \pm 0.002$ | $0.994 \pm 0.002$ |

**Table S4:** SLDB/STRING: Random-walk graph representation learning bias analysis. Explanations and abbreviations as in Table **S2**.

| Model | Evaluation | neg. select | F1 | AUROC | AUPRC | MCC |
|---|---|---|---|---|---|---|
| DeepWalk CBOW | test | Node-degree | $0.830 \pm 0.011$ | $0.898 \pm 0.006$ | $0.866 \pm 0.020$ | $0.676 \pm 0.015$ |
| DeepWalk CBOW | train | Node-degree | $0.911 \pm 0.006$ | $0.955 \pm 0.005$ | $0.909 \pm 0.015$ | $0.819 \pm 0.012$ |
| DeepWalk CBOW | test | Uniform | $0.864 \pm 0.014$ | $0.941 \pm 0.005$ | $0.927 \pm 0.011$ | $0.757 \pm 0.019$ |
| DeepWalk CBOW | train | Uniform | $0.945 \pm 0.007$ | $0.980 \pm 0.003$ | $0.955 \pm 0.008$ | $0.891 \pm 0.013$ |
| DeepWalk GloVe | test | Node-degree | $0.638 \pm 0.012$ | $0.885 \pm 0.001$ | $0.873 \pm 0.001$ | $0.494 \pm 0.009$ |
| DeepWalk GloVe | train | Node-degree | $0.877 \pm 0.005$ | $0.970 \pm 0.000$ | $0.966 \pm 0.000$ | $0.775 \pm 0.007$ |
| DeepWalk GloVe | test | Uniform | $0.592 \pm 0.011$ | $0.797 \pm 0.002$ | $0.713 \pm 0.002$ | $0.339 \pm 0.009$ |
| DeepWalk GloVe | train | Uniform | $0.824 \pm 0.005$ | $0.906 \pm 0.001$ | $0.879 \pm 0.001$ | $0.649 \pm 0.007$ |
| DeepWalk SkipGram | test | Node-degree | $0.878 \pm 0.006$ | $0.921 \pm 0.006$ | $0.875 \pm 0.019$ | $0.756 \pm 0.014$ |
| DeepWalk SkipGram | train | Node-degree | $0.930 \pm 0.008$ | $0.959 \pm 0.006$ | $0.905 \pm 0.018$ | $0.858 \pm 0.016$ |
| DeepWalk SkipGram | test | Uniform | $0.926 \pm 0.007$ | $0.976 \pm 0.002$ | $0.970 \pm 0.006$ | $0.863 \pm 0.011$ |
| DeepWalk SkipGram | train | Uniform | $0.978 \pm 0.005$ | $0.994 \pm 0.002$ | $0.983 \pm 0.005$ | $0.956 \pm 0.009$ |
| First-order LINE | test | Node-degree | $0.860 \pm 0.002$ | $0.930 \pm 0.001$ | $0.935 \pm 0.001$ | $0.710 \pm 0.004$ |
| First-order LINE | train | Node-degree | $0.883 \pm 0.002$ | $0.955 \pm 0.001$ | $0.953 \pm 0.001$ | $0.756 \pm 0.003$ |
| First-order LINE | test | Uniform | $0.922 \pm 0.001$ | $0.968 \pm 0.001$ | $0.972 \pm 0.000$ | $0.848 \pm 0.002$ |
| First-order LINE | train | Uniform | $0.945 \pm 0.001$ | $0.986 \pm 0.000$ | $0.985 \pm 0.000$ | $0.889 \pm 0.001$ |
| HOPE | test | Node-degree | $0.819 \pm 0.004$ | $0.903 \pm 0.003$ | $0.902 \pm 0.009$ | $0.656 \pm 0.009$ |
| HOPE | train | Node-degree | $0.824 \pm 0.004$ | $0.914 \pm 0.003$ | $0.908 \pm 0.009$ | $0.664 \pm 0.010$ |
| HOPE | test | Uniform | $0.869 \pm 0.003$ | $0.945 \pm 0.002$ | $0.959 \pm 0.002$ | $0.779 \pm 0.003$ |
| HOPE | train | Uniform | $0.874 \pm 0.002$ | $0.964 \pm 0.001$ | $0.968 \pm 0.001$ | $0.787 \pm 0.002$ |
| Second-order LINE | test | Node-degree | $0.845 \pm 0.003$ | $0.920 \pm 0.002$ | $0.928 \pm 0.002$ | $0.677 \pm 0.007$ |
| Second-order LINE | train | Node-degree | $0.859 \pm 0.004$ | $0.940 \pm 0.002$ | $0.939 \pm 0.003$ | $0.706 \pm 0.008$ |
| Second-order LINE | test | Uniform | $0.920 \pm 0.001$ | $0.966 \pm 0.001$ | $0.972 \pm 0.001$ | $0.846 \pm 0.002$ |
| Second-order LINE | train | Uniform | $0.934 \pm 0.002$ | $0.983 \pm 0.001$ | $0.983 \pm 0.001$ | $0.871 \pm 0.003$ |
| Walklets CBOW | test | Node-degree | $0.876 \pm 0.022$ | $0.919 \pm 0.007$ | $0.893 \pm 0.017$ | $0.780 \pm 0.027$ |
| Walklets CBOW | train | Node-degree | $0.976 \pm 0.003$ | $0.993 \pm 0.005$ | $0.986 \pm 0.010$ | $0.952 \pm 0.006$ |
| Walklets CBOW | test | Uniform | $0.893 \pm 0.024$ | $0.974 \pm 0.007$ | $0.970 \pm 0.009$ | $0.821 \pm 0.033$ |
| Walklets CBOW | train | Uniform | $0.994 \pm 0.002$ | $0.997 \pm 0.001$ | $0.993 \pm 0.002$ | $0.988 \pm 0.005$ |
| Walklets GloVe | test | Node-degree | $0.541 \pm 0.005$ | $0.500 \pm 0.002$ | $0.499 \pm 0.001$ | $0.001 \pm 0.003$ |
| Walklets GloVe | train | Node-degree | $0.545 \pm 0.005$ | $0.504 \pm 0.001$ | $0.503 \pm 0.001$ | $0.006 \pm 0.002$ |
| Walklets GloVe | test | Uniform | $0.544 \pm 0.005$ | $0.500 \pm 0.002$ | $0.483 \pm 0.001$ | $0.012 \pm 0.003$ |
| Walklets GloVe | train | Uniform | $0.547 \pm 0.005$ | $0.504 \pm 0.001$ | $0.486 \pm 0.001$ | $0.017 \pm 0.002$ |
| Walklets SkipGram | test | Node-degree | $0.819 \pm 0.033$ | $0.914 \pm 0.014$ | $0.883 \pm 0.025$ | $0.703 \pm 0.035$ |
| Walklets SkipGram | train | Node-degree | $0.976 \pm 0.003$ | $0.994 \pm 0.004$ | $0.988 \pm 0.009$ | $0.953 \pm 0.006$ |
| Walklets SkipGram | test | Uniform | $0.839 \pm 0.031$ | $0.984 \pm 0.004$ | $0.977 \pm 0.006$ | $0.751 \pm 0.037$ |
| Walklets SkipGram | train | Uniform | $0.995 \pm 0.002$ | $0.997 \pm 0.001$ | $0.994 \pm 0.002$ | $0.990 \pm 0.004$ |

**Table S5:** STRING: Random-walk graph representation learning bias analysis. Explanations and abbreviations as in Table **S2**.

[2] Luca Cappelletti, Tommaso Fontana, Elena Casiraghi, Vida Ravanmehr, Tiffany J. Callahan, Marcin P. Joachimiak, Christopher J. Mungall, Peter N. Robinson, Justin T. Reese, and Giorgio Valentini. Grape: fast and scalable graph processing and embedding. *CoRR*, abs/2110.06196, 2021.
